# Telomeric repeat distribution and chromosomal evolution in the slipper orchid genus *Paphiopedilum*

**DOI:** 10.64898/2025.12.30.697028

**Authors:** Tianying Lan, Victor A. Albert

## Abstract

The chromosomal mechanisms responsible for the extraordinary diversity in chromosome number and karyotype structure in *Paphiopedilum* remain incompletely resolved, particularly the processes by which extensive dysploid variation has arisen in the apparent absence of polyploidy. Telomeric repeats provide informative cytogenetic markers for reconstructing chromosome evolution because their typically terminal localization and repetitive nature can retain signatures of breakage, fusion, and repair. To address the chromosomal evolutionary problem in *Paphiopedilum*, telomeric repeat distribution was examined in 41 species representing all major taxonomic sections (phylogenetic lineages) using fluorescence in situ hybridization (FISH). In addition to conventional terminal telomeres, diverse interstitial telomeric repeat (ITR) patterns were detected, including centromeric, pericentromeric, and subtelomeric arrays, as well as rare chromosome ends lacking detectable telomeric signals. Classification of these patterns into twelve chromosomal types, interpreted in a phylogenetic context, suggests that telomere-associated and chromosome structural processes have contributed to karyotype diversification, including centric fission, subtelomeric rearrangement, repeat amplification, and potentially centric fusion, with different processes predominating among lineages. In some sections, such as *Pardalopetalum*, these types resolve into internally consistent patterns often dominated by terminal localization with recurrent subtelomeric expansion, providing a clear cytogenetic signature of lineage-specific telomere repeat dynamics, whereas other sections exhibit greater within-section heterogeneity in both signal position and intensity. These results provide a coherent cytogenetic framework for dysploid chromosome evolution in *Paphiopedilum* and identify telomeric repeat dynamics as an important contributor to karyotype diversification in the genus.

## INTRODUCTION

*Paphiopedilum*, together with *Phragmipedium* and *Mexipedium*, comprise the conduplicate, strap-leaved members of the slipper orchid subfamily Cypripedioideae, a lineage long recognized for extraordinary diversity in chromosome number and karyotype structure^1-6^. Early-branching sections of the paleotropical genus *Paphiopedilum* are typically characterized by diploid chromosome numbers of 2*n* = 26 (*x* = 13), whereas more phylogenetically derived sections exhibit a wide range of chromosome numbers forming an extended dysploid series^4^. These patterns are not readily explained by progressive gain or loss of individual chromosomes in a classical aneuploid sense, but instead are most parsimoniously interpreted as the outcome of repeated structural chromosome rearrangements, including centric fission and, possibly, fusion events (cf. ^7^).

Within Cypripedioideae, a phylogenetic framework helps define what is basal versus derived for karyotype traits. Multiple molecular phylogenetic studies support *Mexipedium* as sister to the neotropical genus *Phragmipedium*, with that pair together sister to *Paphiopedilum*^8-16^. Cytological work indicates that *Phragmipedium* also spans a broad dysploid range^3^, but a parallel one based on a different fundamental chromosome number of *x* = 9, with some species exhibiting reduced chromosome numbers dominated by metacentric chromosomes, whereas *Mexipedium* (also neotropical) retains 2*n* = 26 with a distinct karyotype organization^2^. Placing *Paphiopedilum* telomere-pattern evolution in this broader cypripedioid context emphasizes that extensive structural repatterning and dysploid series are not unique to *Paphiopedilum*, but occur across the subfamily.

G. Ledyard Stebbins proposed that dysploid chromosome series can provide a powerful mechanism for speciation by generating partial reproductive barriers through structural heterozygosity^7,17^. In this framework, changes in chromosome number and morphology are filtered by their effects on meiotic pairing and fertility, allowing structurally viable rearrangements to accumulate over time. These ideas are particularly relevant to *Paphiopedilum*, a genus characterized by frequent hybridization, fragmented distributions, and small population sizes^18^.

Because chromosome rearrangements often entail breakage, stabilization, or rejoining of chromosome ends, telomeric DNA emerges as a key molecular component linking structural change to chromosomal function. Telomeric DNA protects chromosome ends and facilitates proper replication and segregation^19^. In most flowering plants, telomeric DNA consists of tandem arrays of the Arabidopsis-type repeat (TTTAGGG)n localized at chromosome termini^20^. However, telomeric repeats are also frequently detected at interstitial chromosomal positions in diverse taxa^19,21-23^. Such interstitial telomeric repeats (ITRs) have often been interpreted as remnants of historical chromosome fusion events, though alternative mechanisms including repeat amplification and repair-associated insertion have also been proposed^24^.

At the molecular level, studies in model plant systems have demonstrated that interstitial telomeric repeats are dynamic genomic elements closely linked to DNA breakage and repair pathways^20,25^. Telomeric repeats can be inserted at double-strand break (DSB) sites through telomerase-mediated healing or nonhomologous end joining, particularly in subtelomeric regions, and may subsequently undergo rapid amplification or dispersal through recombination-mediated processes^24-26^.

Although genomic resources for Cypripedioideae remain limited (but see ^27^), available genome size data^28^ and repeat analyses indicate that slipper orchid genomes are exceptionally repeat-rich^29,30^. This genomic architecture provides a plausible substrate for recurrent breakage, rearrangement, and repeat amplification, suggesting that telomeric repeat dynamics are tightly linked to broader patterns of genome evolution in the group.

In this study, fluorescence in situ hybridization was used to examine telomeric repeat distribution across a broad phylogenetic sampling of *Paphiopedilum*. By integrating cytological observations with phylogenetic relationships, classical chromosomal theory, and emerging molecular perspectives, this work aims to clarify the structural processes underlying dysploid chromosome evolution in this genus.

Comparative cytogenetics and genome sequencing in other angiosperm groups provide important precedents for interpreting interstitial telomeric repeats in *Paphiopedilum*. In the Brassicaceae, comparative chromosome painting and genome-scale synteny have shown that modern karyotypes with reduced chromosome numbers can retain clear signatures of ancestral chromosomes fused into composite chromosomes, and these signatures can be resolved as mosaics of conserved genomic blocks spanning inferred fusion junctions^31-35^. This synthesis of cytology and genomics has been developed in classic Arabidopsis-centered work on chromosome-number reduction and refined by later block-based and phylogenomic analyses across the family, e.g., conserved block systems and protochromosome reconstructions^36,37^.

In the latter Brassicaceae “paleogenomics” analyses, where ancestral chromosomal blocks are tracked through triplicated subgenomes and modern chromosome structures, explicit details on chromosome-scale fusions, translocations, and inversions have been reconstructed with high confidence with chromosome-level resources available^38,39^. Such systems establish a concrete framework for what it would mean, in *Paphiopedilum*, to connect cytological signals (such as ITRs) to molecular breakpoints: one would seek conserved colinearity blocks that traverse or abut candidate junctions, along with repeat landscapes enriched near those junctions.

Outside of such genome-resolved systems, however, comparable inferences about chromosome restructuring must often rely on classical cytogenetic evidence and phylogenetic replication across lineages. Among monocots, Araceae provide a particularly relevant comparative case because dysploid series^40^ and interstitial telomere-like repeats have been examined together in a phylogenetic context^41^. Studies applying telomeric probes across Araceae dysploid lineages^42^ show that internal telomeric signals can occur in association with inferred fusion/fission scenarios, yet may also be absent in some dysploid karyotypes, underscoring that ITRs are not uniquely diagnostic of any single rearrangement pathway and may be lost or remodeled over time.

Within Orchidaceae itself, dysploidy framed explicitly as fusion/fission has been documented in *Heterotaxis*, where chromosome-number variation was evaluated in a phylogenetic framework and interpreted via competing fission and fusion hypotheses^43^. The presence of such a case within orchids (see also ^44-46^) provides direct justification for treating fusion/fission-driven dysploidy as a general orchid mechanism rather than a *Paphiopedilum* idiosyncrasy.

Recent organellar genomic resources for Cypripedioideae also support a view of the group as genomically dynamic. Comparative plastome and mitogenome analyses have revealed substantial differences in plastome size, inverted repeat structure, *ndh* gene presence/absence across genera and species, and evidence for horizontal gene transfer^47-49^. These patterns do not directly determine nuclear karyotype evolution, but they reinforce the broader point that slipper orchid genomes, across compartments, may show atypical structural trajectories.

Finally, models of ITR origin that go beyond simple fusion relics^26,50^ are increasingly supported by a broader literature on telomere and subtelomere dynamics. These considerations motivate explicit mechanistic hypotheses for *Paphiopedilum*, in which subtelomeric and pericentromeric contexts may differ not only in their propensities for rearrangement but also in the likelihood that resulting chromosomal configurations are retained over evolutionary time, given that disruption of centromere function is expected to impose disproportionately severe fitness costs relative to telomere remodeling^51^.

## MATERIALS AND METHODS

Forty-one species of *Paphiopedilum* representing all sections recognized by V.A. Albert^16,52^ were analyzed. Information on the species and sections is provided in **Table 1**. Chromosome preparation, probe labeling, and FISH were accomplished exactly as in Lan and Albert^8^, with the exception that different probe was utilized (*Arabidopsis*-type telomeric DNA was synthesized by PCR using primers (5’-TTTAGGG-3’)5 and (5’-CCCTAAA-3’)5) and a more stringent washing step was used: slides were washed in 2 × SSC at room temperature for 5 min, followed by three times in 2 ×SSC, 0.1% SDS at room temperature for 5 min and three times in 0.1×SSC, 0.1% SDS at 37° C for 5 min.

**Table 1.** *Paphiopedilum* species studied, chromosome numbers, and telomeric repeat distribution types.

| Taxon | Sect. *Concoloria* | Telomere Distribution Types |
| --- | --- | --- |
| *Paphiopedilum* |  |  |
| Subg. *Parvisepalum* |  |  |
| Sect. *Parvisepalum* |  |  |
| *armeniacum* | 26 | I, II, V |
| *delenatii* | 26 | I, IV, V, VI |
| *emersonii* | 26 | I, II, III, V |
| *hangianum* | 26 | I, II, III, V |
| *malipoense* | 26 | I, II, III, V |
| *micranthum* | 26 | I, II, III, V |
| Subg. *Brachypetalum* |  |  |
| Sect. *Concoloria* |  |  |
| *bellatulum* | 26 | I |
| *niveum* | 26 | I |
| Subg. *Paphiopedilum* |  |  |
| Sect. *Cochlopetalum* |  |  |
| *liemianum* | 32 | I |
| *moquetteanum* | 32 | I |
| *primulinum* | 32 | I |
| *victoria-mariae* | 34 | I |
| *victoria-regina* | 34 | I |
| Sect. *Paphiopedilum* |  |  |
| *druryi* | 30 | I |
| *fairrieianum* | 26 | I, VIII, IX |
| *henryanum* | 26+(2B) | I, VII |
| *hirsutissimum* | 26 | I |
| *tigrinum* | 26 | I, IV, V, VIII, X |
| *villosum* | 26 | I |
| Sect. *Coryopedilum* |  |  |
| *adductum* | 26 | I, IX, X, XI, XII |
| *gigantifolium* | 26 | I, IX, X, XI, XII |
| *glanduliferum* | 26 | I |
| *kolopakingii* | 26 | I |
| *randsii* | 26 | I, IX, X, XI, XII |
| *rothschidianum* | 26 | I |
| *sanderianum* | 26 | I |
| *stonei* | 26 | I |
| *supardii* | 26 | I, IX, X, XI, XII |
| Sect. *Pardalopetalum* |  |  |
| *dianthum* | 26 | I, IX, X, XI |
| *haynaldianum* | 26 | I, X |
| *parishii* | 26 | I, IX, X, XI |
| Sect. *Barbata* |  |  |
| *acmodontum* | 38 | I |
| *curtisii* | 36 | I |
| *callosum* | 32 | I |
| *dayanum* | 36 | I |
| *hennisianum* | 34 | I |
| *purpuratum* | 40 | I |
| *sangii* | 28 | I |
| *sukhakulii* | 40 | I |
| *venustum* | 40 | I |
| *wardii* | 42 | I, IX, X |

## RESULTS

### Section *Parvisepalum*

Section *Parvisepalum* (all species of which are 2*n* = 26) represents the sister lineage to all other species of *Paphiopedilum*. All six species of *Parvisepalum* studied show a considerable number of interstitial telomeric repeat (ITR) signals, including both small dot-like signals and strong band-like signals (**Figure 1**). Most of the dot-like signals were found in centromeric regions, whereas band-like signals were located in pericentromeric regions, the latter in some cases almost covering the entire chromosome arms. The most extensive distribution was observed in *Paphiopedilum micranthum*, which has four band-like signals and numerous ITRs on every chromosome (**Figure 1A**). Similar patterns were observed in *P. malipoense*, *P. emersonii*, and *P. hangianum*, but two to four chromosomes in these species lacked ITRs altogether. However, *P. armeniacum* shows many fewer ITRs and only two band-like signals. Interestingly, in *P. delenatii* there are twelve chromosomes harboring band-like signals, but no dot-like ITR signals were observed.

**Figure 1.**
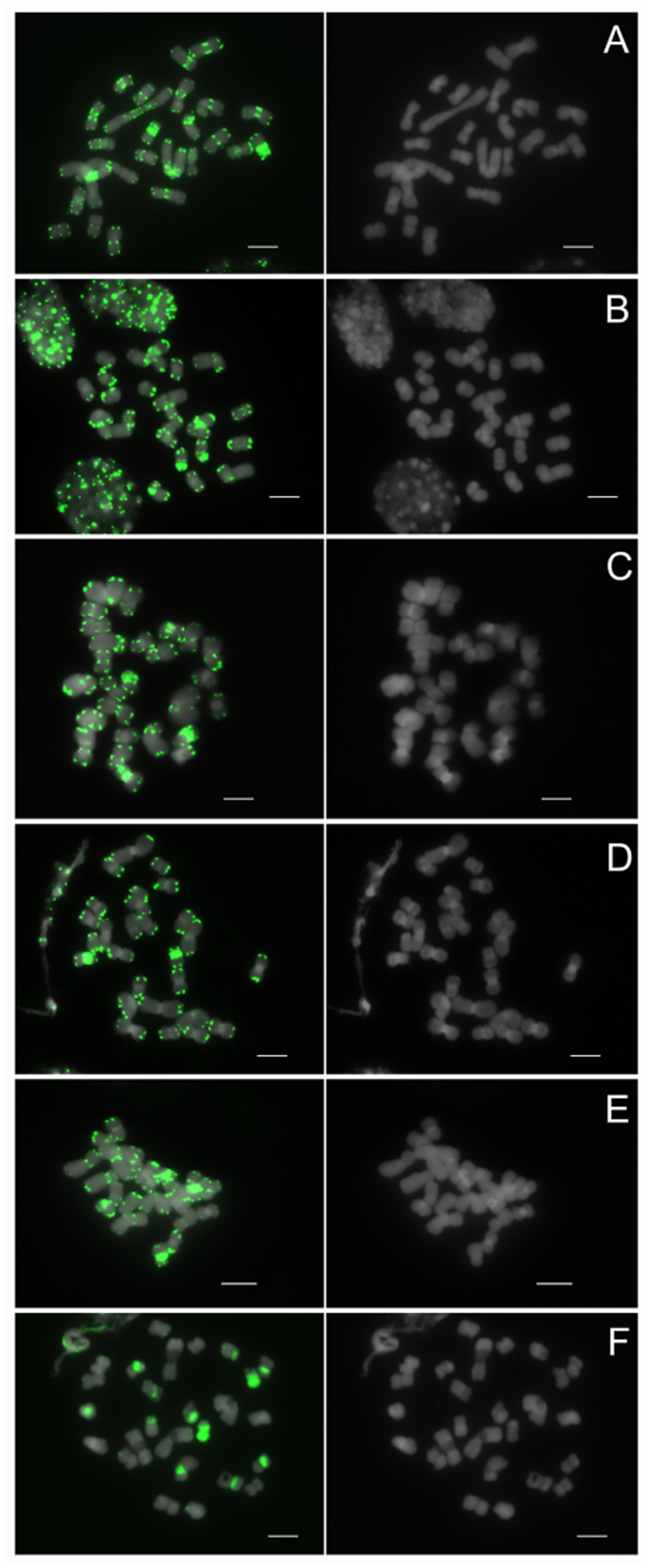
Fluorescence in situ hybridization (FISH) distribution of telomeric sequences on metaphase chromosomes of *Paphiopedilum* section *Parvisepalum*. (**A**) *Paphiopedilum micranthum*, (**B**) *P. malipoense*, (**C**) *P. emersonii*, (**D**) *P. armeniacum*, (**E**) *P. hangianum*, (**F**) *P. delenatii*. Telomeric sequences (green) were detected using FISH; chromosomes were counterstained with DAPI (gray). Scale bars = 10 µm.

### Section *Concoloria*

Based on current phylogenetic information, after the branching off of *Parvisepalum*, section *Concoloria* (all 2*n* = 26) is the sister group of the remaining *Paphiopedilum* species. Both species of section *Concoloria* studied, *P. niveum* and *P. bellatulum*, display dot-like signals on the chromosome termini, representing a conventional telomere localization pattern, and no ITRs were observed.

### Section *Cochlopetalum*

As in section *Concoloria*, in section *Cochlopetalum* (where a dysploid series occurs^4^), only conventional telomere localization patterns were observed in all five species studied (**Figure 3**).

**Figure 2.**
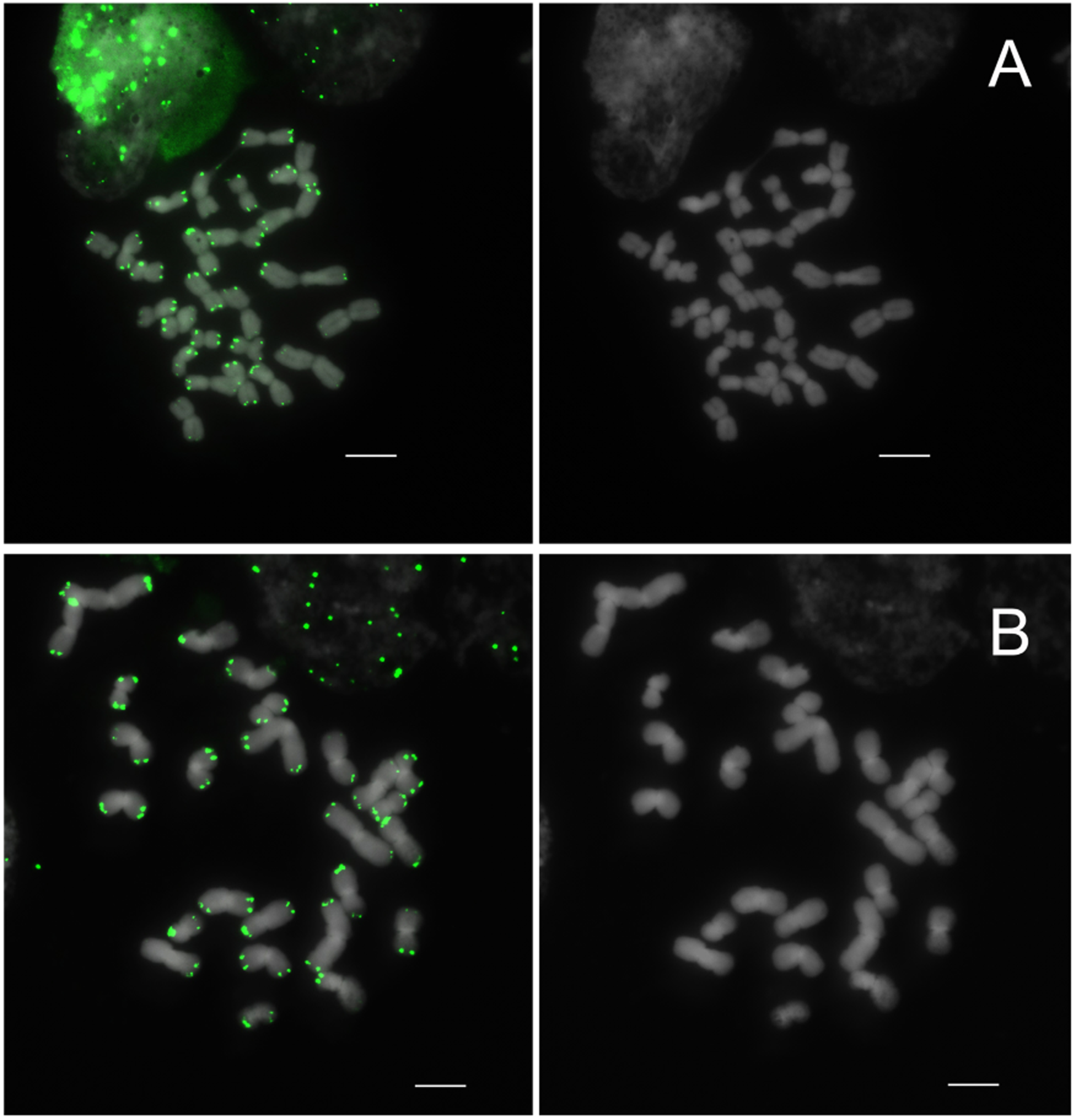
FISH distribution of telomeric sequences on metaphase chromosomes of *Paphiopedilum* section *Concoloria*. (**A**) *Paphiopedilum niveum*, (**B**) *P. bellatulum*. Telomeric sequences are shown in green; chromosomes were counterstained with DAPI (gray). Scale bars = 10 µm.

**Figure 3.**
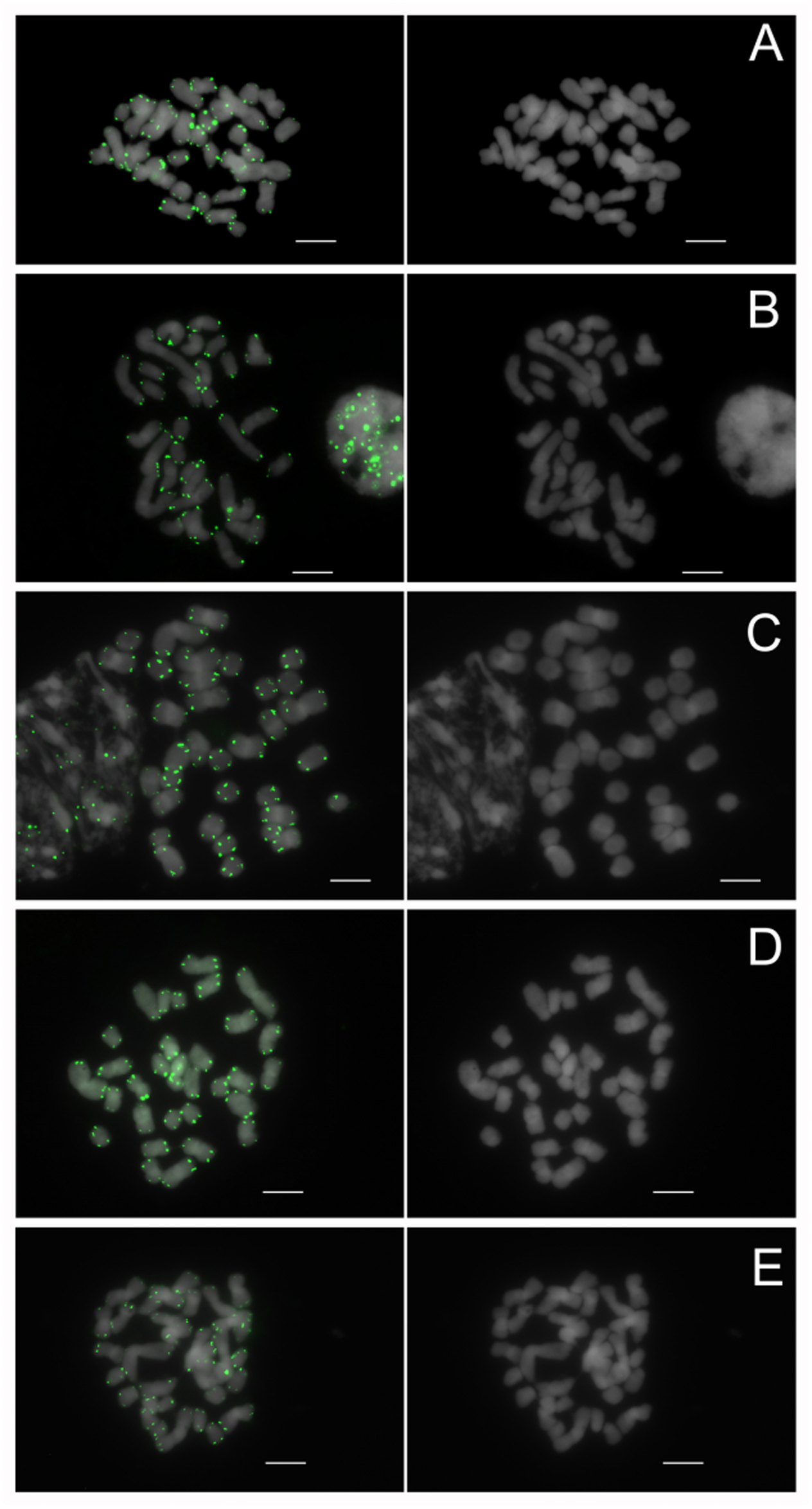
FISH distribution of telomeric sequences on metaphase chromosomes of *Paphiopedilum* section *Cochlopetalum*. (**A**) *Paphiopedilum liemianum*, (**B**) *P. moquettianum*, (**C**) *P. victoria-regina*, (**D**) *P. primulinum*, (**E**) *P. victoria-mariae*. Telomeric sequences are shown in green; chromosomes were counterstained with DAPI (gray). Scale bars = 10 µm.

### Section *Paphiopedilum*

Diverse telomeric distribution patterns were observed in section *Paphiopedilum* (all 2*n* = 26) (**Figure 4**). In *Paphiopedilum druryi*, *P. hirsutissimum*, and *P. villosum*, no ectopic telomeric signals were detected. In contrast, in *P. tigrinum*, fourteen band-like signals (one band on each of fourteen chromosomes) were observed, located either in subtelomeric or pericentromeric regions. In *P. fairrieanum*, four weak band-like signals were detected in subtelomeric regions. Notably, a breakage event resulting in broken chromosome ends lacking detectable telomeric repeats was observed in *P. henryanum* (**Figure 4A**, arrowheads); this state was observed in four independent cells.

**Figure 4.**
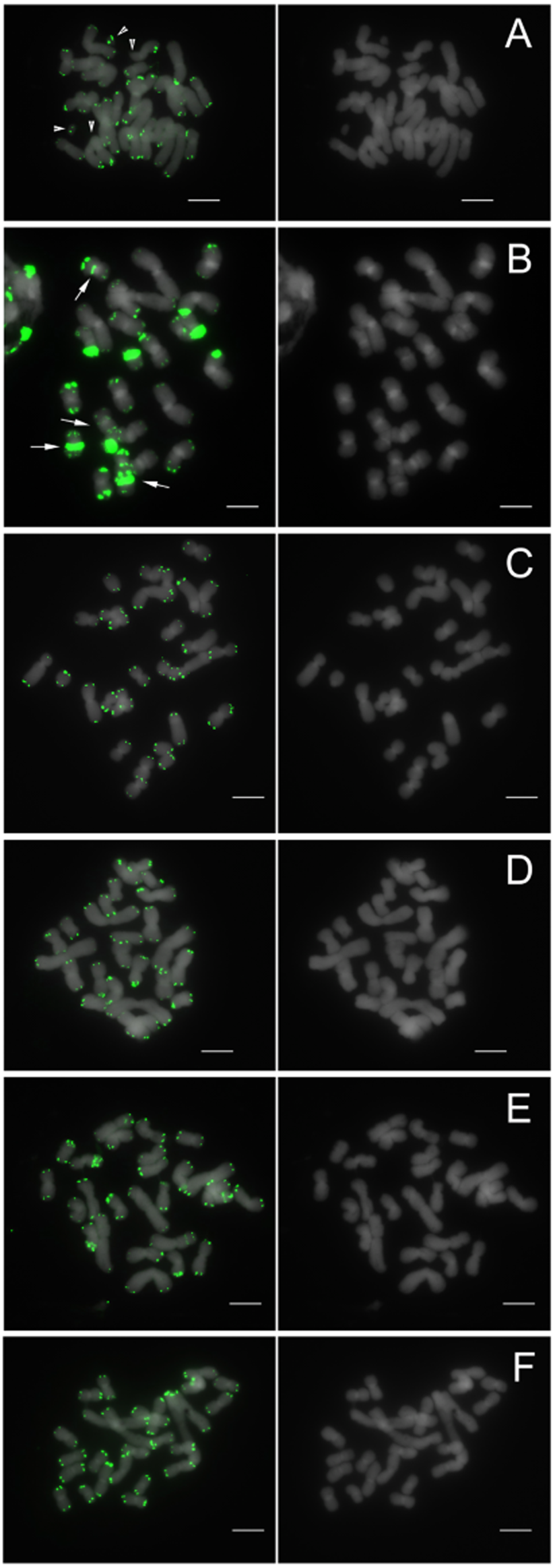
FISH distribution of telomeric sequences on metaphase chromosomes of *Paphiopedilum* section *Paphiopedilum*. (**A**) *Paphiopedilum henryanum*, (**B**) *P. tigrinum*, (**C**) *P. druryi*, (**D**) *P. hirsutissimum*, (**E**) *P. fairrieanum*, (**F**) *P. villosum*. Arrows indicate pericentromeric or centromeric interstitial telomeric repeats (ITRs) in *P. tigrinum*. Arrowheads indicate chromosome ends lacking detectable telomeric signals in *P. henryanum*. Scale bars = 10 µm.

### Sections *Coryopedilum* and *Pardalopetalum*

In sections *Coryopedilum* (**Figure 5**) and *Pardalopetalum* (all species of which are 2*n* = 26) (**Figure 6**), where telomeric repeat positions are even more diverse than in section *Paphiopedilum*, the conventional telomere distribution pattern was observed in five species, whereas numerous strong band-like signals, all located in telomeric and subtelomeric regions, were present in the remaining seven species studied. The number of chromosomes harboring ectopic signals varies from eight in *Paphiopedilum haynaldianum* to sixteen in *P. dianthum*, *P. adductum*, and *P. supardii*.

**Figure 5.**
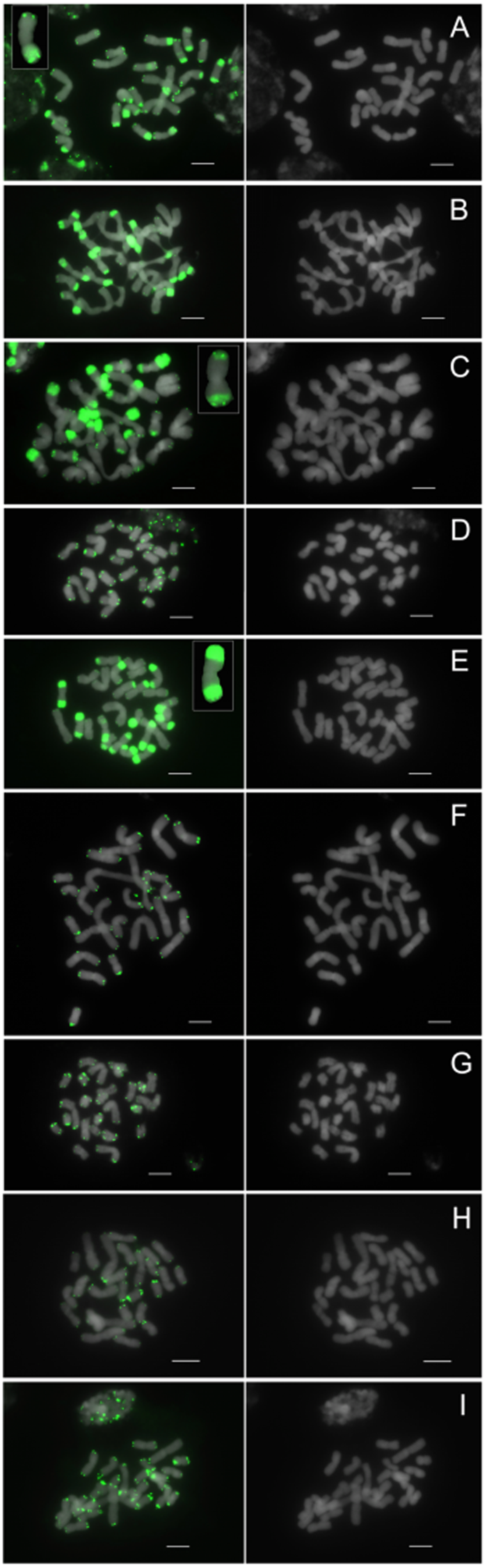
FISH distribution of telomeric sequences on metaphase chromosomes of *Paphiopedilum* section *Coryopedilum*. (**A**) *Paphiopedilum randsii*, (**B**) *P. gigantifolium*, (**C**) *P. adductum*, (**D**) *P. stonei*, (**E**) *P. supardii*, (**F**) *P. glanduliferum*, (**G**) *P. sanderianum*, (**H**) *P. kolopakingii*, (**I**) *P. rothschildianum*. Enlarged chromosomes in (**A**), (**C**), and (**E**) show representative chromosomes bearing physically separated subtelomeric telomeric bands and functional terminal telomeres. Scale bars = 10 µm.

**Figure 6.**
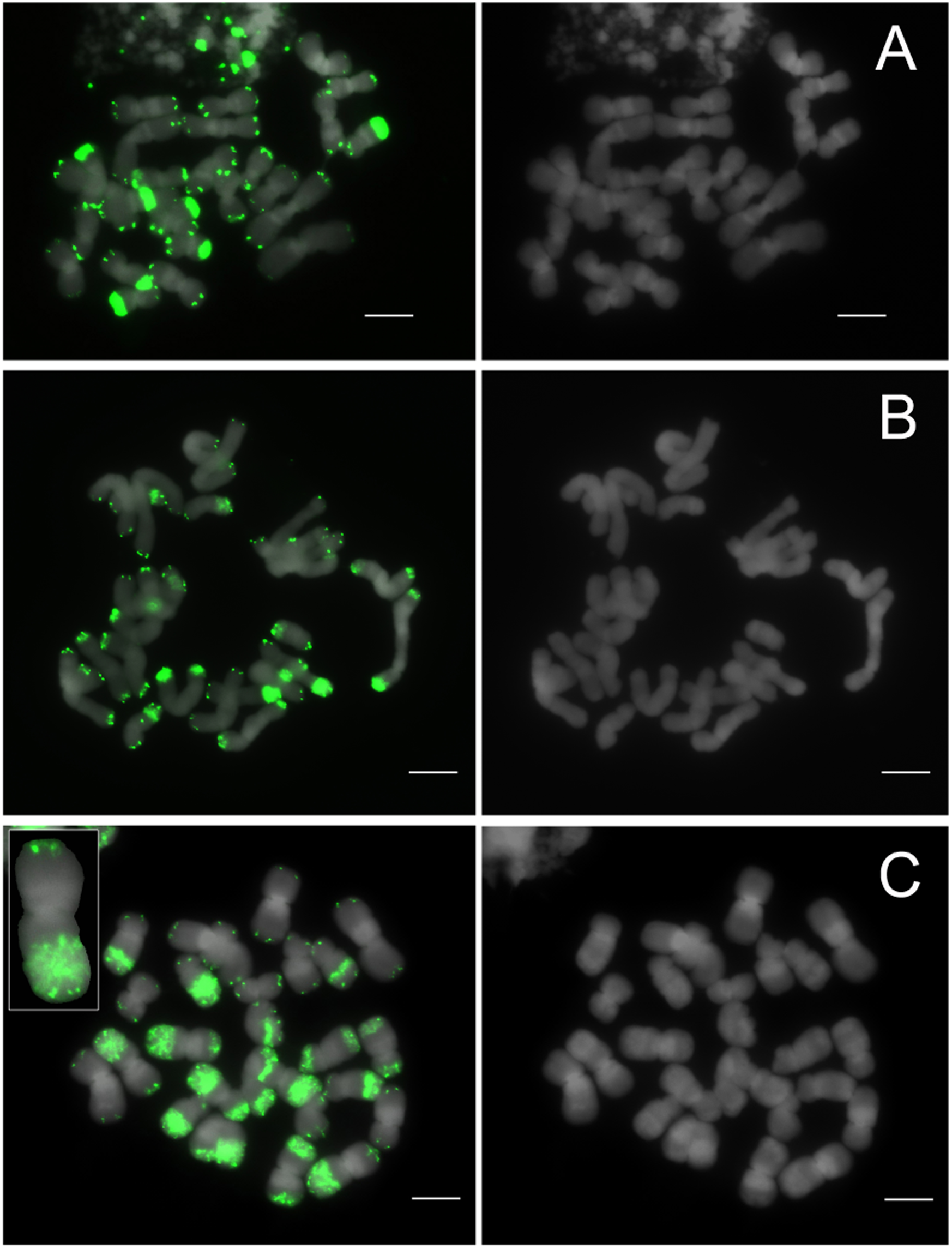
FISH distribution of telomeric sequences on metaphase chromosomes of *Paphiopedilum* section *Pardalopetalum*. (**A**) *Paphiopedilum haynaldianum*, (**B**) *P. parishii*, (**C**) *P. dianthum*. The enlarged chromosome in (**C**) illustrates a representative chromosome bearing physically separated subtelomeric telomeric bands and functional terminal telomeres. Scale bars = 10 µm.

### Section *Barbata*

All species in section *Barbata* (where an dysploid series occurs^4^), except for *Paphiopedilum wardii*, exhibit the conventional terminal localization pattern (**Figure 7**). In *P. wardii*, eight chromosomes harbor strong band-like signals in telomeric and subtelomeric regions.

**Figure 7.**
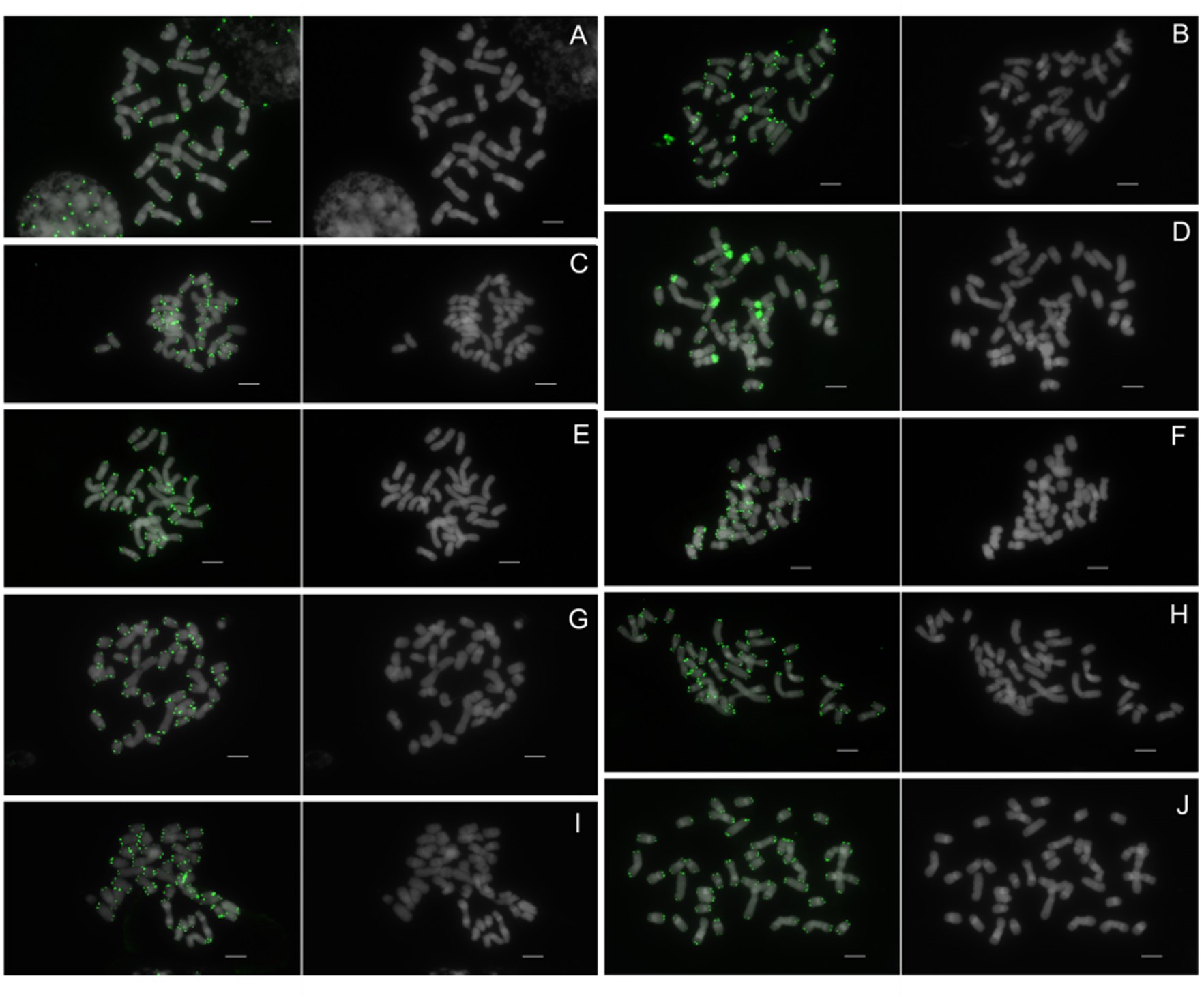
FISH distribution of telomeric sequences on metaphase chromosomes of *Paphiopedilum* section *Barbata*. (**A**) *Paphiopedilum sangii*, (**B**) *P. curtisii*, (**C**) *P. purpuratum*, (**D**) *P. wardii*, (**E**) *P. callosum*, (**F**) *P. hennisianum*, (**G**) *P. dayanum*, (**H**) *P. acmodontum*, (**I**) *P. venustum*, (**J**) *P. sukhakulii*. Telomeric sequences are shown in green; chromosomes were counterstained with DAPI (gray). Scale bars = 10 µm.

**Figure 8.**
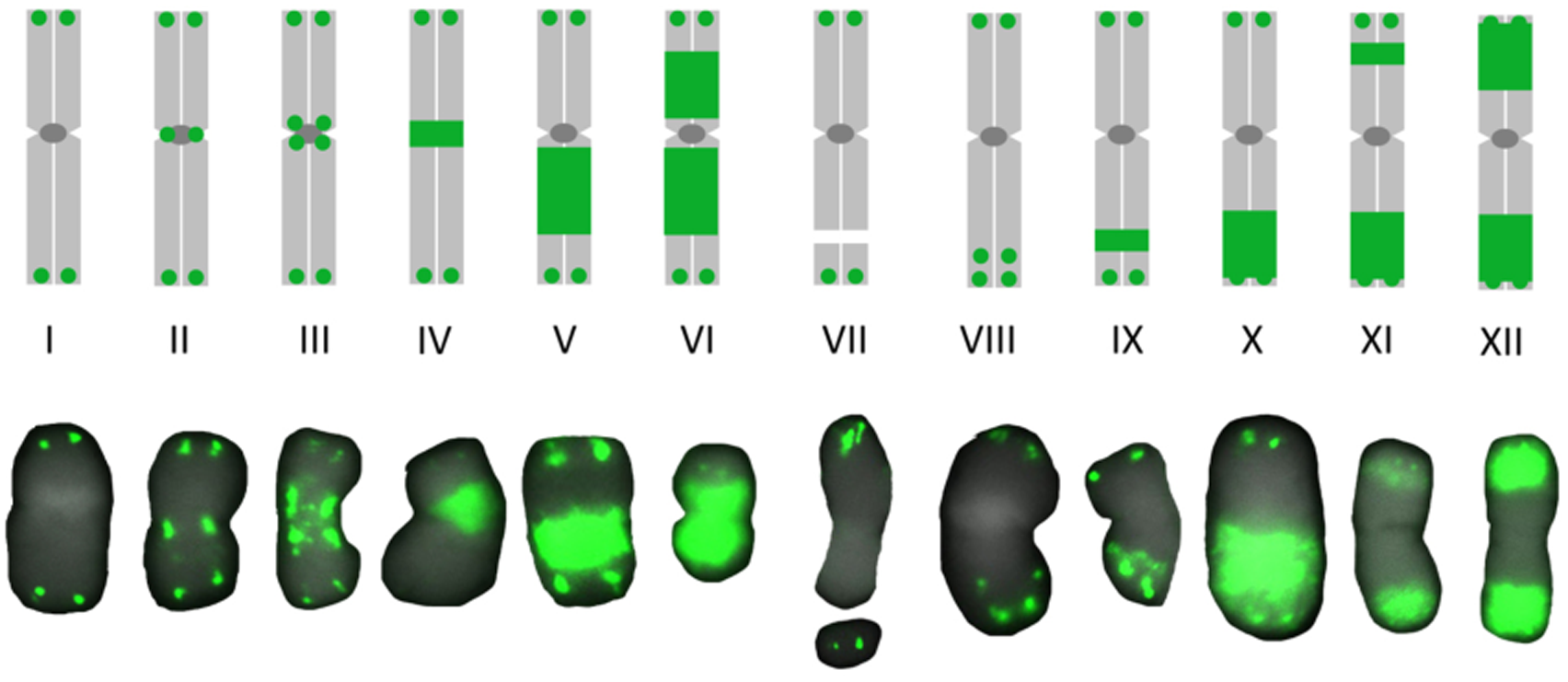
and a model for the evolution of *Paphiopedilum* telomeric repeat patterns. Classification of twelve chromosome types based on telomeric repeat distribution patterns:

- **Type I**, conventional chromosomes with dot-like telomeric signals at both ends
- **Type II**, a single dot-like telomeric signal at the centromeric region of each sister chromatid
- **Type III**, two or more discrete dot-like signals in centromeric and pericentromeric regions
- **Type IV**, a band-like signal at the centromeric region
- **Type V**, a strong band-like signal at the pericentromeric region
- **Type VI**, two strong band-like signals in the pericentromeric regions of both arms
- **Type VII**, absence of detectable telomeric signals at chromosome ends
- **Type VIII**, dot-like signals in the subtelomeric region
- **Type IX**, a small band-like signal in the subtelomeric region
- **Type X**, a strong band-like signal in the subtelomeric region of one chromosome arm
- **Type XI**, a small band-like signal at one end and a strong band-like signal at the opposite end
- **Type XII**, strong band-like signals in the subtelomeric regions of both arms.

### Classification of distribution patterns into twelve types

According to the positions and relative signal strength observed in the FISH experiments, the general distribution patterns of telomeric repeats are classified into twelve chromosomal types (**Figure 8**). The types exhibited in the various species are summarized in **Table 1**.

## DISCUSSION

The present study reveals extensive diversity in telomeric repeat distribution across *Paphiopedilum* and demonstrates that these patterns are tightly structured by phylogeny and karyotype history. When interpreted in the broader context of Cypripedioideae evolution and classical chromosomal theory, the results highlight the role of repeated chromosome structural rearrangements in shaping genomic architecture.

From a karyotypic perspective, these phylogenetically structured telomeric patterns align with long-standing models of chromosome number evolution in the genus. *Paphiopedilum* exhibits increasing chromosome number with phylogenetic progression while largely conserving total chromosome arm number. These patterns are consistent with centric fission as a major driver of karyotype evolution, but the telomeric repeat data suggest that centric fusion, subtelomeric translocations, inversions, and deletions may also contribute substantially. Genome size dynamism known in *Paphiopedilum* ^2,28^ suggests the possibility that amplification of interstitial telomeric repeats might contribute to genome expansion through the formation of new telomeres or centromeres, as in potato^22^.

Interpreting these cytological patterns requires distinguishing between telomeric sequence presence and telomere function. Functional versus non-functional telomeric arrays are a central interpretive issue for *Paphiopedilum* because telomeric probes detect sequence identity rather than telomere function. In plants, functional telomeres are typically constrained to the chromosome termini and, while they may vary in length, they rarely extend to encompass megabase-scale subtelomeric domains^19,20,53,54^. Consequently, the very large, band-like signals that span substantial portions of terminal chromosome regions in some *Paphiopedilum* species are most parsimoniously interpreted as expanded telomere-like repeat arrays embedded within subtelomeric DNA rather than intact telomere tracts that directly serve end-protection roles. This distinction matters mechanistically: non-functional telomere-like arrays can behave as recombinogenic repeats that promote ectopic exchanges^24,50,54,55^ while true telomeres are generally protected by specialized chromatin and telomere-binding proteins that suppress end-to-end fusions and inappropriate repair^20^.

When viewed in a phylogenetic context, this distinction between functional and non-functional telomeric arrays becomes especially informative. The phylogenetic co-occurence of species having abundant interstitial telomeric repeats with closely related species lacking such signals implies that telomeric repeat amplification and dispersal can proceed rapidly once initiated. Differences between pericentromeric and subtelomeric interstitial repeat distributions suggest strong chromosomal context effects, with subtelomeric regions known to undergo frequent nonhomologous end joining-mediated translocations and subsequent homologous recombination^19,24,54-56^. These mechanisms provide a plausible explanation for the phylogenetically radiative spread of subtelomeric interstitial telomeric repeats observed in derived *Paphiopedilum* sections.

Notably, this subtelomeric-biased pattern is not universal across the genus. In the early-branching *Parvisepalum* lineage, strong ITR bands are often few in number and concentrated near centromeric/pericentromeric regions. A parsimonious explanation is a survivorship filter: rearrangements and unequal exchanges in centromere-adjacent regions are more likely to compromise centromere function and be eliminated, whereas analogous events in subtelomeric regions can be tolerated provided chromosome ends remain capped or are efficiently re-capped. This lineage-specific contrast mirrors broader principles of chromosome survivorship observed across eukaryotes.

Subtelomeric regions are widely recognized among diverse phyla as fast-evolving domains marked by segmental duplication, ectopic recombination, and frequent sequence exchange among nonhomologous chromosome ends. In Arabidopsis, subtelomeric regions show evidence of complex similarity patterns consistent with recurrent nonhomologous exchanges, providing a model for how telomere-proximal repeats can spread without requiring full chromosome fusions^57^. More generally, reviews of telomere and subtelomere dynamics emphasize that somatic double-strand break repair in plants and other eukaryotes often proceeds via non-homologous end joining (NHEJ), while homologous recombination (HR) can act on duplicated segments and repeat arrays^58-60^; the interplay of these pathways provides multiple routes by which telomere-like repeats can be inserted, amplified, and redistributed.

These mechanistic considerations reinforce an important interpretive caution: interstitial telomeric repeats can be historical markers of rearrangements, but they are not uniquely diagnostic of any single event type. Depending on genomic context and time since formation, ITRs may be eroded, dispersed, or rendered cytologically undetectable. Thus, the absence of ITRs in a lineage (for example, in *Paphiopedilum* sections showing exclusively terminal localization) does not necessarily imply the absence of historical rearrangements; it can also reflect loss, remodeling, or replacement of repeats after the events occurred. These patterns are consistent with comparative cytogenetic and genomic studies in other plant lineages, which show that dysploidy and internal telomeric sequences may be variably coupled or decoupled depending on rearrangement history and lineage context.

Within this interpretive framework, *Paphiopedilum* provides an opportunity to move from inference to explicit hypothesis testing. The patterns documented here invite integration with broader concepts of repeat-mediated genome restructuring, including the possibility that ITRs themselves influence chromosome behavior^50^. In *Paphiopedilum*, this leads to a testable hypothesis that chromosomes bearing abundant ITRs might exhibit elevated local rearrangement rates or altered meiotic recombination landscapes relative to chromosomes lacking such arrays, which could feed back into karyotype diversification.

A parallel case of repeat instability in *Paphiopedilum* reinforces this perspective. The dynamic behavior of telomeric repeats in the genus resembles previously documented instability of ribosomal DNA loci in *Paphiopedilum*^8,61^. This relationship warrants more extensive consideration because both represent major repeat families with distinct but potentially interacting evolutionary trajectories. rDNA arrays are well-known to undergo rapid copy-number change and positional turnover^62^, and rDNA loci can act as chromosomal fragile sites or breakage-prone domains under some conditions^63-65^. The observation that some species show both dispersed rDNA and strong ITRs, while others do not, suggests partial coupling: shared upstream drivers (e.g., elevated breakage or repeat turnover) may promote both phenomena, but telomere-repeat amplification also requires telomere-specific pathways and subtelomeric exchange processes to reach the extreme ITR distributions seen in multiple derived *Paphiopedilum* sections.

Genome size evolution provides a complementary, integrative axis for evaluating these processes. Massive amplification of ITRs, like amplification of other tandem repeats, can be expected to contribute to genome expansion, and species with exceptionally abundant ITRs plausibly accumulate substantial repeat mass in subtelomeric domains. At the same time, centric fission can increase the number of chromosome ends and may be associated with the creation of new telomeric tracts at healed breaks; depending on how breaks are processed, fission could also promote local repeat turnover and expansion. These ideas predict that genome size, chromosome number, rDNA abundance, and ITR burden need not be tightly linked in a single monotonic trend, but can instead show section- and lineage-specific correlations depending on the prevailing restructuring routes. Such decoupling sets the stage for considering how chromosomal restructuring feeds into evolutionary divergence.

Incorporating Stebbins’ perspective ^7^ more explicitly, *Paphiopedilum* becomes a candidate system in which dysploid series and chromosomal repatterning may contribute to speciation by creating structural heterozygosity and reducing gene flow between karyotypic variants. This possibility is strengthened by the genus’ ecological and evolutionary context. Natural hybridization both ancient and recent is broadly documented in *Paphiopedilum*^13,14,66-68^, and hybridization between lineages with differing chromosome architectures can expose meiotic incompatibilities that impose strong selection for karyotype stabilization or for rearrangements that improve pairing. Meanwhile, many species are local endemics with fragmented distributions (cf. ^18^), conditions under which drift and local selection^69^ can facilitate the fixation of rearrangements that might be purged in large, well-mixed populations^70,71^.

Testing these ideas directly is now becoming feasible as genomic resources mature. In Cypripedioideae, emerging genomic datasets provide concrete opportunities for integration, including expanded comparative cytogenetic analyses of *Phragmipedium* and the generation of chromosome-level nuclear assemblies to test whether interstitial telomeric repeats in *Paphiopedilum* coincide with rearrangement junctions, conserved synteny breakpoints, centromere relics, or subtelomeric duplication mosaics. Ongoing genome projects focused on *Paphiopedilum gigantifolium* ^72^, *Phragmipedium kovachii*^73^, and *Mexipedium xerophyticum*^15^ – undertaken by the senior author and collaborators – are well positioned to address these questions.

In light of our findings and discussion, we propose a model for the evolution of the twelve types of telomeric patterns observed in *Paphiopedilum*:

### Type I — Conventional terminal telomeres (baseline/stabilized)

**Cytological pattern:** dot-like signals confined to both chromosome ends.

**Mechanistic interpretation:** either (i) ancestral/stable state with no major telomere-repeat redistribution, or (ii) a post-rearrangement state where internal tracts never formed, were too small to detect, or were erased by turnover. While type I can be an ancestral state, it may also be an “endpoint” after clean healing or after erosion of internal tracts.

Type I therefore does not exclude past rearrangement; it simply indicates absence of detectable internal telomeric-repeat accumulation at the time of observation.

### Type II — Single centromeric dot

**Cytological pattern:** one discrete dot-like signal at centromere (often one per sister chromatid regionally).

**Mechanistic implication:** a localized, limited insertion or retention event at/near the centromere-proximal break site, with minimal subsequent amplification. This is the simplest “repair-associated insertion” signature: one seed tract persists without expanding into a band. Under a fission-only scenario, Type II is the most parsimonious footprint of a single misrouted repair/healing episode at a centric break.

### Type III — Multiple centromeric/pericentromeric dots

**Cytological pattern:** two or more discrete dot-like signals in centromeric and pericentromeric regions.

**Mechanistic implication:** recurrence. Either multiple independent repair-associated insertions occurred (multiple DSBs, repeated fission/breakage episodes), or a small seed tract dispersed locally within pericentromeric repeat arrays (short-range duplication and movement within heterochromatin). Importantly, Type III differs from Type II mainly by multiplicity. It signals that the centromere-proximal environment is repeatedly producing or retaining telomeric motifs, consistent with a breakage-prone, repair-active, repeat-dynamic centromeric domain.

### Type IV — Centromeric band

**Cytological pattern:** a band-like telomeric-repeat signal centered on the centromere. **Mechanistic implication:** amplification of a centromeric seed. Type IV is best explained by repeat expansion within the core centromere repeat landscape. Once a telomeric motif is present, unequal exchange, replication slippage, or break-induced replication (BIR) - like processes can enlarge it from “dot” to “band.” The centromere-core localization suggests amplification constrained to the kinetochore-adjacent domain or to a centromere-associated satellite array incorporating telomeric motifs.

### Type V — Pericentromeric band

**Cytological pattern:** a strong band-like signal in pericentromeric region, often offset from the exact centromere.

**Mechanistic implication:** amplification in flanking heterochromatin. Pericentromeres are frequently more permissive to large repeat expansion than the functional centromere core, and they may experience distinct replication/repair dynamics. Type V therefore suggests a seed tract that either formed in the pericentromere or migrated/expanded preferentially into pericentromeric heterochromatin. It is consistent with a repair-associated insertion near the centromere followed by biased expansion into the pericentromeric neighborhood.

### Type VI — Symmetric pericentromeric bands on both arms

**Cytological pattern:** two strong bands in pericentromeric regions, one on each arm, often roughly symmetric.

**Mechanistic implication:** a shared initiating context plus duplication or bilateral expansion. Symmetry is the key clue: it is hard to explain by two fully independent insertions of identical magnitude in mirror positions. More plausible routes include (i) amplification from a centromere-adjacent seed that expands bidirectionally into both pericentromeric flanks, (ii) duplication of pericentromeric repeat blocks during repair (e.g., template switching, BIR, segmental duplication), or (iii) centromere repositioning and subsequent amplification that yields paired flanking bands. Type VI therefore maps to “amplification-with-structure,” not simply “more amplification.”

### Type VII — No signal at broken ends

**Cytological pattern:** broken chromosome ends lacking detectable telomeric repeats. **Mechanistic implication:** an instability snapshot. Type VII is crucial because it demonstrates that telomere addition is not automatic. It implies recent/ongoing breakage where ends have not yet been capped by sufficient telomeric repeats to be detected, or where ends are degraded/processed in a way that eliminates detectable tracts. It may represent (i) a transient intermediate prior to healing, (ii) a failure mode where broken ends persist, or (iii) rapid cycles of breakage and repair that repeatedly reset ends before stable telomere formation. Type VII may derive directly from delayed/misrouted repair because it is conceptually a “failed or incomplete stabilization” outcome rather than a stable amplified state.

### Type VIII — Subtelomeric dots

**Cytological pattern:** dot-like signals located in subtelomeric region (near ends but not strictly terminal).

**Mechanistic implication:** subtelomeric seeding with minimal expansion. This can arise when a telomeric-repeat tract is introduced near the end via rearrangement (e.g., an internal tract moved toward the end by inversion/translocation) or by repair-associated insertion at a subtelomeric DSB. Because subtelomeres are dynamic, even small seeds can persist; Type VIII indicates the presence of a seed without major local amplification (yet).

### Type IX — Small subtelomeric band

**Cytological pattern:** small band-like signal in subtelomeric region.

**Mechanistic implication:** modest local amplification of a subtelomeric seed. Subtelomeres can amplify repeats rapidly through unequal exchange among duplicated segments and interchromosomal recombination networks. Type IX suggests that amplification has begun but has not yet produced large, intense bands.

### Type X — Strong subtelomeric band on one arm

**Cytological pattern:** strong band-like signal in subtelomeric region of one chromosome arm.

**Mechanistic implication:** asymmetric amplification and/or arm-specific history. The one-arm restriction is informative: it implies (i) a seed that is present on only one end, (ii) differential amplification rates between ends, or (iii) a rearrangement history that relocated or created a seed at only one end. Type X is a robust signature of subtelomeric amplification that is not genome-wide uniform but contingent on local architecture and recombination context.

### Type XI — Mixed ends

**Cytological pattern:** one end shows a small band, the other shows a strong band. **Mechanistic implication:** temporal or mechanistic asymmetry between ends. The mixed pattern is best explained by stepwise histories: one end experienced earlier seeding/amplification (strong band), while the other has a later or weaker seeding event (small band). Alternatively, both ends were seeded but only one end entered a high-amplification regime due to local subtelomeric duplication networks. Type XI therefore encodes “non-synchronous end histories” within a single chromosome.

### Type XII — Strong subtelomeric bands on both arms

**Cytological pattern:** strong band-like signals at subtelomeric regions of each arm. **Mechanistic implication:** dual-end seeding plus parallel amplification. Type XII can arise if both ends carry telomeric-repeat seeds (through redistribution, multiple subtelomeric breaks, or end-to-end exchange across chromosomes) and both ends undergo substantial amplification. In subtelomeric domains, interchromosomal exchange can synchronize repeat landscapes across ends, so Type XII is consistent with a “network effect” where duplicated subtelomeric segments and recombination facilitate parallel amplification at both chromosome ends.

